# The *C. elegans* proteasome subunit RPN-12 is required for hermaphrodite germline sex determination and oocyte quality

**DOI:** 10.1101/2020.06.11.146472

**Authors:** Lourds M. Fernando, Jeandele Elliot, Anna K. Allen

**Affiliations:** Department of Biology Howard University, Washington, DC 20059, USA

**Keywords:** RPN-12, proteasome, *C. elegans*, oogenesis, sex determination

## Abstract

**Background:** The proteasome is a multi-subunit complex and a major proteolytic machinery in cells. Most subunits are essential for proteasome function, and depletion of individual subunits normally results in lethality. RPN-12/Rpn12/PSMD8 is a lid subunit of the 19S regulatory particle (RP) of the 26S proteasome. Studies in *Caenorhabditis elegans* demonstrated that RNAi depletion of RPN-12 does not result in lethality. RPN-12 has not been well studied in higher eukaryotes. In this study we investigate the biological significance of RPN-12 in *C. elegans.*

**Results:** We found that the null mutant *rpn-12(av93)* did not cause major impairment of the proteolytic activity of the proteasome. Most *rpn-12(av93)* hermaphrodites lack sperm leading to feminization of the germ line that can be partially rescued by mating to males. The lack of sperm phenotype can be suppressed by downregulation of TRA-1, a player in the hermaphrodite germline sex determination pathway. Also, *rpn-12(av93)* animals show significant nuclear accumulation of the meiotic kinase WEE-1.3, a protein predominantly localized to the perinuclear region. Interestingly, chemical inhibition of the proteasome did not cause nuclear accumulation of WEE-1.3.

**Conclusions:** RPN-12 plays a previously unknown role in oogenesis and the germline sex determination pathway in *C. elegans* hermaphrodites.

## Introduction

The 26S proteasome is a large protein complex that plays an essential role in regulating protein homeostasis in cells, and is crucial for the development and survival of all organisms.^1–3^ Dysfunction of the proteasome has been linked to lethality, heart disease, aging, and neurodegenerative diseases such as Alzheimer’s and Huntington’s.^4–7^ Hyperactivation of the proteasome is observed in many cancers and chemical proteasome inhibitors are used to treat certain cancers such as multiple myeloma.^7–9^ Canonically, the proteasome acts with the small molecule ubiquitin to comprise the Ubiquitin Proteasome System (UPS).^3^ The UPS marks proteins to be degraded with ubiquitin and then shuttles the ubiquitinated proteins to the proteasome where they are broken down into smaller peptides and the amino acids recycled.^1,3,10^ Alternative pathways by which proteins are targeted to the proteasome have been uncovered, such as monoubiquitination and ubiquitin independent pathways.^11,12^ Additionally, there is scientific precedent in the literature showing that the proteasome has the ability to perform non-proteolytic functions in cell cycle regulation, mRNA export, transcription, heterochromatin spreading and immunity in various organisms.^13–16^

Structural and functional studies of the 26S proteasome have shown that it is composed of different protein subunits elegantly arranged into two 19S regulatory particles (RP) capping a 20S core particle (CP).^17–19^ These subunits are highly conserved from yeast to mammals, and historically the unicellular yeast model systems *Saccharomyces cerevisiae* and *Saccharomyces pombe* have been utilized to elucidate the various roles of the proteasome subunits.^20^ This includes the identification of specific subcomplexes that the proteasomal subunits form, and how these subcomplexes are configured and spatiotemporally distributed in various subcellular compartments.^17,18,21^ For instance, Boehringer et al. demonstrated that one 19S RP proteasome subunit (Rpn12) is needed for the incorporation of another subunit (Rpn10) into the fully functional proteasome.^20^

Similar to other eukaryotes, *C. elegans* possess the conserved canonical structural components of the 26S proteasome. The 20S CP is a cylindrical structure comprised of rings formed by seven alpha and seven beta subunits stacked on top of one another (two outer alpha and two inner beta rings).^10,22^ The 20S CP proteasome cannot perform its function *in vivo* without the aid of the 19S RP whose role is to recognize, deubiquitinate, and translocate ubiquitinated proteins into the core particle for degradation^23^. The 19S RP is divided into a lid and a base component.^10,21,22^ The 19S base is composed of six ATPase subunits (RPT-1, RPT-2, RPT-3, RPT-4, RPT-5, and RPT-6) and four non-ATPases (RPN-1, RPN-2, RPN-10 and RPN-13).^22^ The 19S lid particle is comprised of nine subunits: RPN-3, RPN-5, RPN-6.1, RPN-7, RPN-8, RPN-9, RPN-11, RPN-12 and RPN-15 (DSS-1).^22^

Downregulation of most *C. elegans* 19S RP subunits via RNAi individually causes embryonic lethality therefore impeding any further investigation into the role of individual subunits *in vivo.^24^* Downregulation or mutation of RPN-10, RPN-12 or RPN-15/DSS-1 however, does not cause embryonic lethality in *C. elegans.^24–27^* It has been reported though that RNAi knockdown of RPN-12 and RPN-10 together causes embryonic lethality, implying that these two subunits can compensate for each other.^24^ RPN-15/DSS-1 plays an essential role in *C. elegans* oogenesis and *dss-1(tm370)* loss-of-function deletion mutants exhibit sterility due to oogenesis defects.^27^ RPN-10 has an essential role in *C. elegans* hermaphrodite reproduction by regulating the germline sex determination pathway.^26^ Loss-of-function *rpn-10* mutants possess a feminized germ line and are sterile due to this lack of sperm.^26^ RPN-10 is one of the ubiquitin receptors of the proteasome that is involved in degradation of specific substrates through an ubiquitin-interacting motif (UIM) domain.^26^ Meanwhile, the role of RPN-12 in *C. elegans* remains understudied. RPN-12 contains a conserved PCI domain (Proteasome, Cop9 signalosome, eukaryotic Initiation factor 3) which is known to play roles in scaffolding protein complex subunits together and stabilizing protein-protein interactions.^28,29^ However, the particular function of RPN-12 in the proteasome remains unclear.

*C. elegans* serves as a great metazoan model to study the proteasome dynamics. As a multicellular model, it can help elucidate tissue specific roles of proteasome subunits. We are interested in studying the roles of proteasome subunits in the *C. elegans* germ line. The *C. elegans* hermaphrodite germ line is composed of two U-shaped gonad arms connected to a shared uterus. An adult hermaphrodite gonad arm consists of a distal region where the germ cell nuclei are in a syncytium and a proximal region where fully cellularized oocytes line up awaiting ovulation (Figure 1B). At ovulation, an oocyte is pushed into the spermatheca, the sperm-storage structure of the animal, and fertilization occurs before the newly fertilized oocyte is released into the uterus where subsequent embryogenesis occurs (Figure 1B).

**Figure 1.**
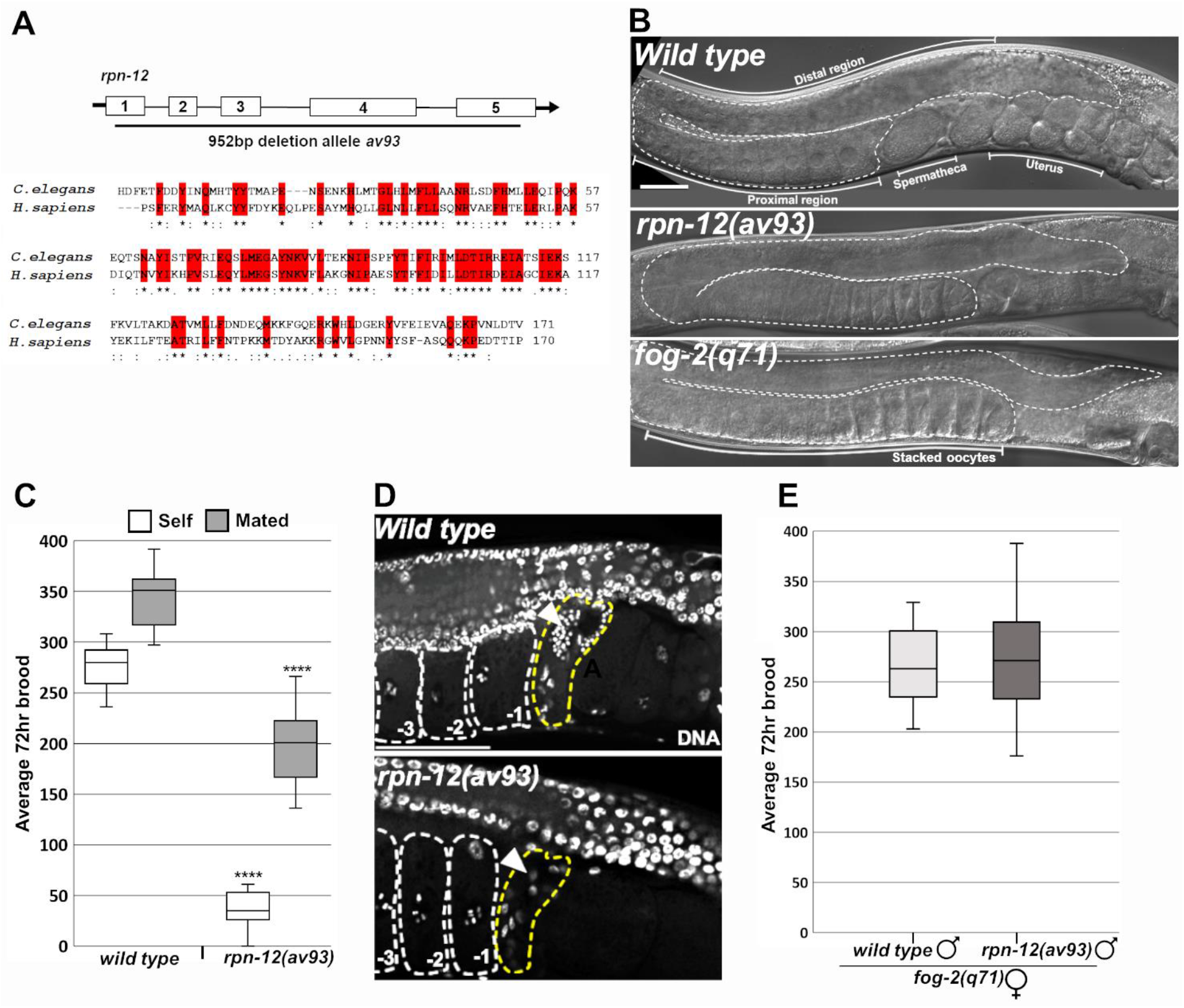
Loss of the conserved C. *elegans* proteasomal subunit RPN-12 causes germline feminization in hermaphrodites. A. Top: Schematic of *rpn-12* containing 5 exons (boxes). The *av93* mutation, black line below schematic, deletes most of *rpn-12* genomic region (952bp). Bottom: PCI domain alignment of *C. elegans* RPN-12 with *Homo sapiens* PSMD8 showing 42% identity. Red boxes and asterisks (*) indicate identical residues, colons (:) residues with strongly similar properties and periods (.) residues with weakly similar properties. B. DIC images of wild type, *rpn-12(av93)* and *fog-2(q71)* hermaphrodite germ lines. One gonad arm is outlined in white dashes. Wild type germ line labeled: distal region, proximal region, spermatheca and the uterus. Stacked oocytes labeled in *fog-2(q71)* germ line. C. Average 72hour brood of self-fertile (n=26) and cross fertilized (n=28) *rpn-12(av93)* hermaphrodites. Cross-fertilized progeny obtained by mating with wild type males. D. DAPI stained germ lines showing spermatheca (outlined in yellow) in wild type and *rpn-12(av93)* hermaphrodites. Oocytes are outlined in white and −1, −2, and −3 oocytes are labeled. Arrows point to where sperm should reside. E. Average 72hr brood of wild type and *rpn-12(av93)* males mated to *fog-2(q71)* females. Error bars indicate maximum and minimum values. Medians represented by line within box. ****p <0.0001 as determined by Student *T-*test. Scale bars indicate 50μm.

Here we report a role for RPN-12 in the *C. elegans* germ line. Loss of *rpn-12* results in feminized animals possessing otherwise normal viability and lifespan, but reduced oocyte quality. Moreover, we show that RPN-12 is required for the proper oocyte subcellular localization of the *C. elegans* Myt1/Wee1 meiotic kinase, WEE-1.3 and that absence of RPN-12 can suppress a penetrant *wee-1.3(RNAi)* infertility phenotype.^30,31^ Interestingly, chemical inhibition of the proteasome and overall decrease of the proteolytic activity in the animals does not show the same results. Therefore, our results demonstrate that RPN-12 is required for proper germline sex determination and oogenesis, and we speculate that RPN-12 may play a non-proteolytic role in *C. elegans* oocytes through its interactions with WEE-1.3.

## Results

### *rpn-12(av93)* mutants lack sperm

In *C. elegans,* mutation or RNAi depletion of most proteasome subunits individually causes severe proteasome dysfunction and results in embryonic lethality.^24^ This embryonic lethal phenotype precludes further investigation of tissue-specific roles of specific proteasome subunits that may be related or unrelated to a proteolytic function. Interestingly, certain *C. elegans* subunits, such as RPN-10, RPN-12 and DSS-1 (RPN-15), do not cause embryonic lethality when depleted.^24,26,27^ The roles of RPN-10 and DSS-1 in spermatogenesis and oogenesis respectively are previously reported but no analysis of the RPN-12 deletion phenotype has been studied.^26,27^ We generated a strain bearing a 952bp deletion of the coding region of *rpn-12* [AG343-*rpn-12(av93)];* these animals are homozygous viable and show no defects in either larval growth or life span (Figure 1A).^25^

Initial observations suggested that *rpn-12(av93)* animals might have a decreased brood compared to wild type animals. In conducting individual brood size assays only 36% of the *rpn-12(av93)* hermaphrodites were self-fertile at 20°C (data not shown) and the average self brood of those fertile animals was significantly lower when compared to wild type hermaphrodite self brood (Figure 1C). The *rpn-12(av93)* hermaphrodites can be cross fertilized by sperm from wild type males such that 100% of the hermaphrodites are now fertile (data not shown). However, the average brood size of the *rpn-12(av93)* animals mated with wild type males is still moderately but significantly lower than that of the wild type mated hermaphrodites (Figure 1C, compare mated genotypes, p-value = 1.12414e^-06^). The inability of mated *rpn-12(av93)* hermaphrodites to reach a wild-type level of fertility even when sperm is provided suggest that these animals are producing poor quality oocytes.

The fact that mating restored fertility led us to hypothesize that lack of self sperm was contributing to the infertility phenotype. In DIC images of a single wild type gonad arm, the distal gonad region contains numerous germ cells in a syncytium and individualized oocytes form in the proximal gonad arm near the site of sperm storage (spermatheca) (Figure 1B, wild type). Additionally, the uterus of a wild type animal is filled with developing, multi-cellular embryos. DIC images of young adult hermaphrodite germ lines of *rpn-12(av93)* mutants revealed a stacked oocyte phenotype in the proximal gonad arm similar to that exhibited by the *fog-2(q71)* feminized germ line mutants (Figure 1B). Feminized mutants such as these can result from a lack of sperm. We found that while spermathecae of wild type adult hermaphrodites were filled with spermatids, the spermathecae of *rpn-12(av93)* adult hermaphrodites were void of sperm (Figure 1D). Interestingly, male *rpn-12(av93)* animals mated to feminized *fog-2(q761)* animals exhibit no difference in brood size and those males contain on average the same amount of spermatids per gonad area as observed in wild type males (Figure 1E and data not shown). This demonstrates that the sperm defect in *rpn-12(av93)* animals is specific to the process of hermaphrodite spermatogenesis.

### *rpn-12(av93)* mutants exhibit a minor impairment of the proteolytic activity of the proteasome

Proper proteolytic function of the proteasome is important for gametogenesis in *C. elegans.^32^* To assess the extent by which overall proteolytic activity is affected in the *rpn-12(av93)* mutant, we obtained a recently published strain that provides a readout for proteasome dysfunction.^32^ This transgenic strain has a non-hydrolysable ubiquitin molecule that is mutated and fused to a GFP tagged Histone H2B and driven by the germline *pie-1* promoter (UbG76V::GFP::H2B). The mutation in the ubiquitin causes the GFP and H2B to undergo continuous proteasomal degradation in the germ line. Wildtype *C. elegans* hermaphrodites containing this reporter have no GFP expression in the distal gonad and a low amount of GFP expression in the proximal oocyte nuclei (Figure 2A). Animals mutant for *rpn-12* and possessing the UbG76V::GFP::H2B reporter exhibit a slight increase in GFP levels in the distal gonad and the proximal oocytes (Figure 2A-B). This is in contrast to RNAi downregulation of other essential proteasome subunits, such as RPN-6.1 and RPN-7, which exhibit very bright expression of GFP throughout the germ line, indicative of severe proteasome dysfunction upon knockdown of those specific subunits (Figure 2A-B). We can conclude from this that the absence of RPN-12 results in minor inhibition of the proteasome’s proteolytic activity, and importantly this reduction is not significant enough to affect the viability of the *rpn-12(av93)* mutants.

**Figure 2.**
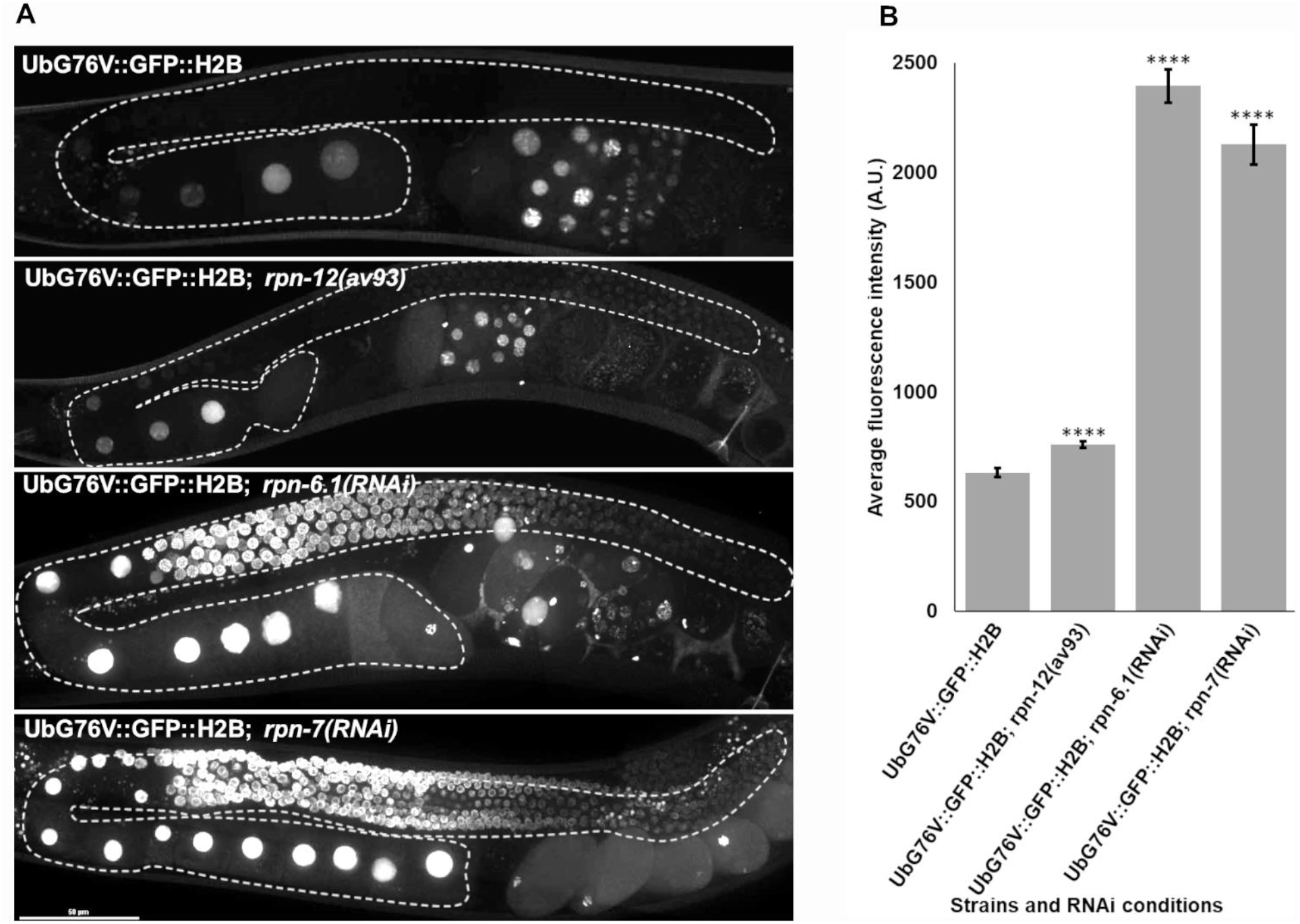
Loss of RPN-12 causes minor impairment of the proteolytic activity of the proteasome. A. Live imaging of gonads containing the UbG76V::GFP::H2B transgene. Animals have either just the transgene, transgene and the *rpn-12(av93)* mutation, or the transgene plus RNAi depletion of either *rpn-6.1* or *rpn-7.* n-values are: UbG76V::GFP::H2B (n=14), UbG76V::GFP::H2B; *rpn-12(av93)* (n=29), UbG76V::GFP::H2B; *rpn-6.1(RNAi)* (n=10) and UbG76V::GFP::H2B; *rpn-7(RNAi)* (n=7). A single gonad arm is outlined in white dashes in each image. B. Average intensity (arbitrary units) of the entire outlined area of the germ lines in A. All experiments were conducted at least three times. ****p <0.0001 determined by Student’s *T-*test. Errors bars represent ± SEM. Scale bars indicate 50μm.

### RPN-12 is ubiquitously expressed in the germ line and soma of *C. elegans*

To better understand the role of RPN-12 in *C. elegans* reproduction, we endogenously tagged the *rpn-12* locus at the N–terminus with GFP via CRISPR. We observe that RPN-12 is expressed ubiquitously throughout both the germ line and soma in *C. elegans* hermaphrodites (Figure 3A). This expression is both cytoplasmic and nuclear, which is clearly seen in the distal germ cells and proximal oocyte nuclei (Figure 3A’-A”). We can also observe RPN-12 expression in the spermatids stored in the spermatheca (Figure 3A’”). The nuclear localization of RPN-12 is similar to the localization of various other proteasome subunits in the 19S RP that our laboratory has tagged via CRISPR (unpublished data). Additionally, RPN-12 colocalizes in the nuclei of proximal oocytes and distal germ cells with a general 20S CP antibody specific to multiple alpha subunits (Figure 3B-C’”). This confirms that the GFP::RPN-12 is localizing to the proteasome which is nuclear and cytoplasmically localized. The insets in Figure 3C-C’” represent one pachytene region nucleus from the distal region that shows both RPN-12 and the 20S CP antibody localized adjacent to the chromatin similar to what has been observed with other published fluorescently tagged proteasome subunits.^32^

**Figure 3.**
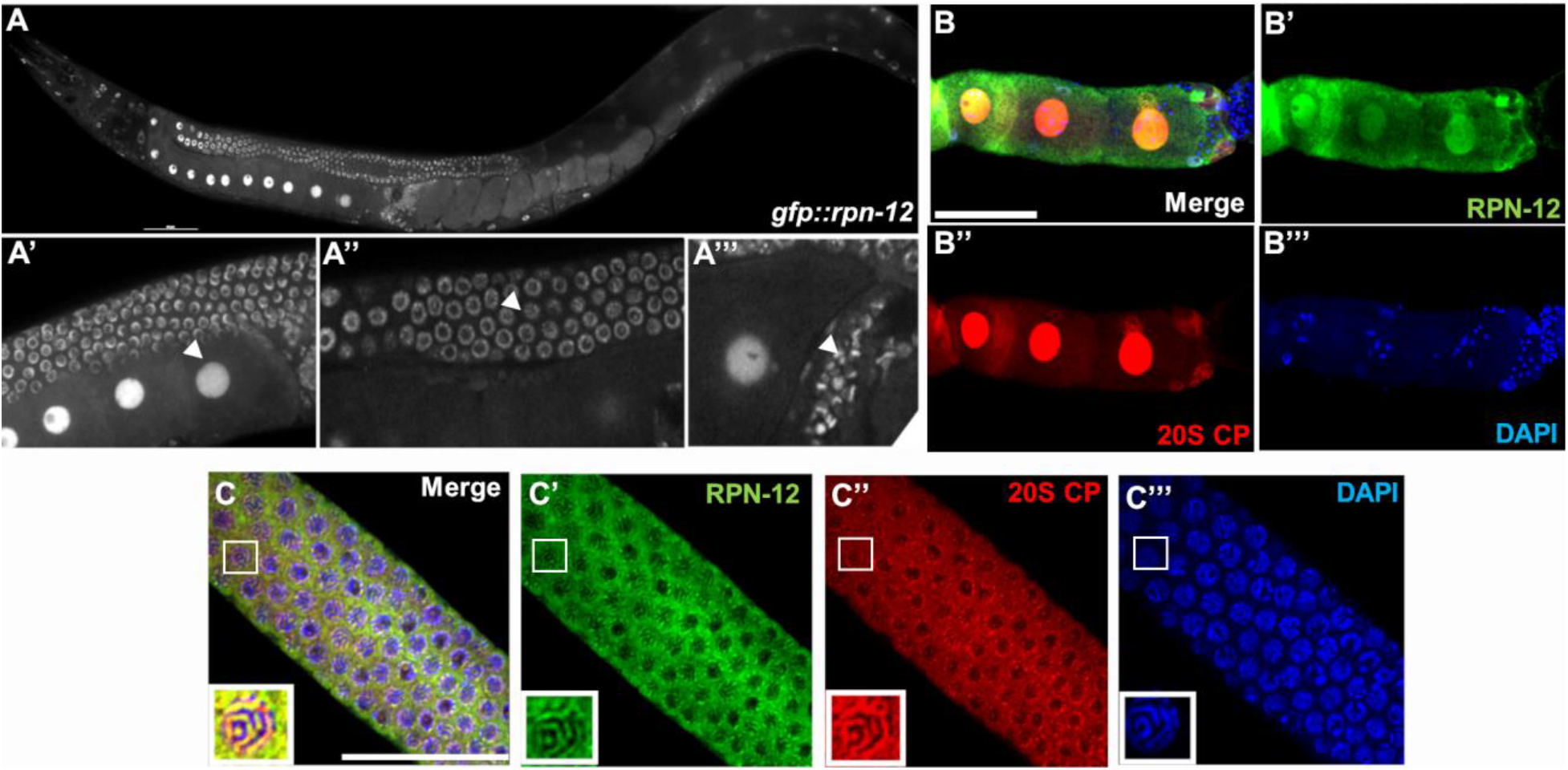
RPN-12 is ubiquitously expressed in the germ line and soma. A. Live image of a hermaphrodite expressing an endogenously GFP tagged RPN-12, *gfp::rpn-12(ana6).* A’ Proximal oocytes showing bright nuclear (white arrowhead) and faint cytoplasmic GFP expression. A” Distal syncytial germ cell nuclei expressing GFP (white arrowhead). A”’ GFP expression in hermaphrodite self sperm (white arrowhead). B. Merged confocal images of the dissected proximal gonad region from *gfp::rpn-12(ana6)* animal immunostained with anti-GFP (green), anti-alpha 20S CP subunits (red) and DAPI (blue). Individual images of GFP::RPN-12 in B’, anti-alpha 20S CP subunits in B” and DNA in B”’. C. Immunostaining of the dissected distal region of *gfp::rpn-12(ana6)* animals stained for GFP (green and C’), the 20S CP alpha subunits (red and C”), and DNA (blue and C”’). The insets show staining in a single pachytene nucleus. Stainings were conducted at least three independent times and over 26 animals were imaged to confirm staining pattern. Scale bars indicate 5Oμm.

### Loss of RPN-12 alters WEE-1.3 localization and suppresses *wee-1.3(RNAi)* infertility

WEE-1.3 is a major *C. elegans* meiotic kinase that belongs to the larger MYT1 family of kinases. It maintains oocyte meiotic arrest in *C. elegans* by providing inhibitory phosphorylations to the CDK-1 component of Maturation Promoting Factor (MPF), thereby inactivating MPF and halting meiosis.^30^ Downregulation of WEE-1.3 via RNAi causes precocious oocyte meiotic maturation, ultimately resulting in the production of fertilization incompetent oocytes and infertile hermaphrodites.^30,31^ In wild type hermaphrodites WEE-1.3 is localized in the cytoplasm, ER and in the perinuclear region of both germ line and somatic cells (Figure 4A).^31^ Interestingly, we observed that WEE-1.3 has an aberrant nuclear localization and accumulation in hermaphrodite oocytes when we utilized RNAi to knockdown specific proteasome subunits, including *rpn-12(RNAi)* (data not shown). To test whether complete loss of RPN-12 causes the same phenotype, we crossed endogenously tagged *gfp::wee-1.3* animals to the *rpn-12(av93)* strain to generate *gfp::wee-1.3; rpn-12(av93)* animals, and observed similar aberrant localization of WEE-1.3 (Figure 4B).

**Figure 4.**
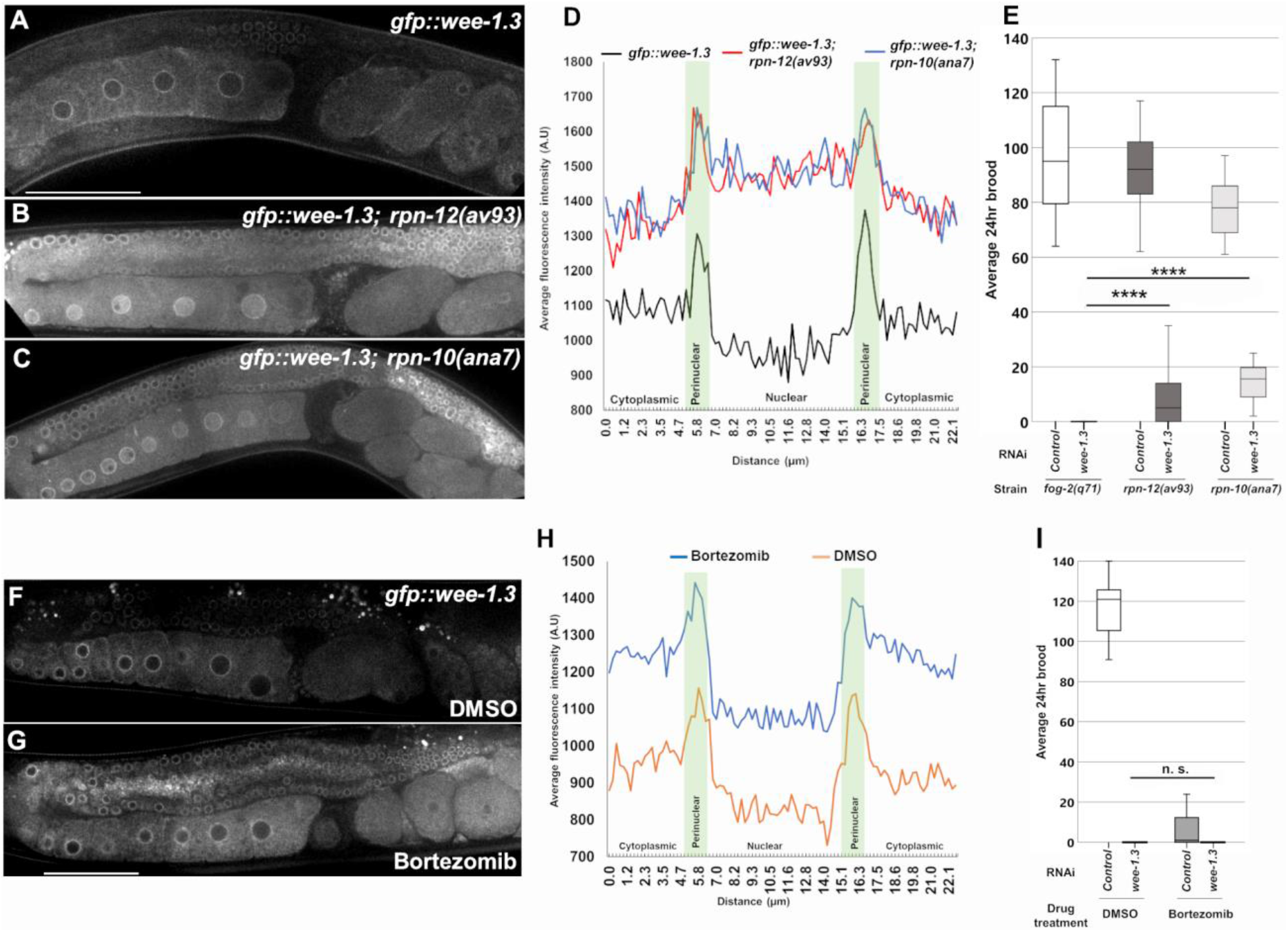
Loss of RPN-12 causes WEE-1.3 accumulation in oocyte nuclei and suppresses *wee-1.3(RNAi)* infertility. Live confocal images of GFP expression in *gfp::wee-1.3* (A)*, gfp::wee-1.3; rpn-12(av93)* (B) and *gfp::wee-1.3; rpn-10(ana7)* (C) animals. All images were taken with the same microscope settings. D. Intensity graph showing GFP expression levels across the −2 oocyte in the cytoplasmic, perinuclear and nuclear regions in the following strains: *gfp::wee-1.3* (black line)*, gfp::wee-1.3; rpn-12(av93)* (red line) and *gfp::wee-1.3; rpn-10(ana7)* (blue line). E. Average 24hr brood of *control(RNAi)* and *wee-1.3(RNAi)* animals in *fog-2(q71)* (n=25–30)*, rpn-12(av93)* (n=23-43) and *rpn-10(ana7)* (n=11-20) strains mated to wild type males as a source of sperm. F-G. Live confocal images of GFP expression in DMSO treated (F) or 100μM Bortezomib treated *gfp::wee-1.3* animals (G). H. Intensity graph of GFP expression across the −2 oocytes in DMSO *gfp::wee-1.3* (red line) and Bortezomib *gfp::wee-1.3* (blue line) treated animals. Perinuclear regions in the intensity graphs shaded in green. I. Average 24hr brood of wild type hermaphrodites treated with either *control(RNAi)* and *wee-1.3(RNAi)* conditions and either DMSO (n=24-30) or 100μM Bortezomib (n=32-42). Medians represented by lines within boxes. Errors bars represent minimum and maximum values. All experiments were conducted at least three times. ****p <0.0001 determined by Student’s *T-*test. Scale bars indicate 50μm.

We were interested in whether this alteration in endogenous WEE-1.3 expression would also be observed in another viable proteasome mutant. Previously, Shimida et al. reported the characterization of an *rpn-10* deletion allele.^26^ This allele was no longer available, so we generated our own CRISPR *rpn-10* deletion mutant *[rpn-10(ana7)]* (see methods for construction of mutant). A similar nuclear accumulation of GFP::WEE-1.3 was observed in *gfp::wee-1.3*; *rpn-10(ana7)* hermaphrodites (Figure 4C).

To quantify the levels of GFP::WEE-1.3, we determined the intensity of fluorescence in the −2 oocyte across the nucleus in the three different strains. Animals only containing endogenous GFP::WEE-1.3 exhibit decreased fluorescence of WEE-1.3 in the nucleus compared to the cytoplasm, with peak expression observed in the perinuclear region (Figure 4D, black line). In contrast, the GFP::WEE-1.3 fluorescence in animals mutant for *rpn-12* or *rpn-*10 demonstrates higher intensity within the nuclei than in the cytoplasm (Figure 4D, red and blue lines). There are still however two distinct peaks of fluorescence intensity that corresponds to the perinuclear region (Figure 4D, green shaded region). We also observed that overall, the levels of WEE-1.3 are higher in the *rpn-12* and *rpn-10* null mutants than in wild type animals (Figure 4D). It is possible that the misexpression of WEE-1.3 in *rpn-12(av93)* animals leads to poor quality oocytes, thereby affecting the reproductive capacity of the animals.

One possible explanation for the overall increase in WEE-1.3 levels could be the slight decrease in proteolytic activity of the proteasome in *rpn-12(av93)* and *rpn-10(ana7)* mutants (Figure 2). This could result in a decrease in WEE-1.3 turnover thereby causing an elevation of apparent WEE-1.3 levels. Interestingly, when we treated *gfp::wee-1.3* animals with a high concentration of the chemical proteasome inhibitor bortezomib (100μM), we did not observe a strong nuclear accumulation of WEE-1.3 compared to DMSO treated animals (Figure 4F-G). As an indication that bortezomib is causing significant loss of proteolytic activity of the proteasome we looked in the uteri of the animals for either dead or developing embryos. Reduced proteolytic activity of the proteasome causes embryonic lethality as seen in down regulation of essential proteasome subunits such as RPN-6.1 or RPN-7.^10,24,33^ Bortezomib treated hermaphrodites all contained dead embryos with abnormal cellular morphology compared to healthy embryos in vehicle treated animals’ uteri (Figure 4F-G). This indicates that the bortezomib is functioning and decreasing the proteasome activity, and that a significant reduction of proteasome activity is not sufficient to result in the aberrant nuclear localization of WEE-1.3.

Quantifying the levels of WEE-1.3 in the −2 oocyte, we observed overall higher levels of WEE-1.3 intensity in bortezomib treated animals supportive of a decrease functionality of the proteasome and thus slow degradation of the GFP::WEE-1.3. Interestingly, however the WEE-1.3 expression pattern in bortezomib treated animals follows that of control treated animals, with higher cytoplasmic WEE-1.3 expression and lower nuclear WEE-1.3 expression (Figure 4H). Taken together, this data suggests that it is not overall proteasome dysfunction that results in the increased nuclear accumulation of WEE-1.3 but rather that this alteration is specific to the absence of either the RPN-12 or RPN-10 proteins.

A previous study by the lab showed that 44 genes when co-depleted with WEE-1.3 via RNAi suppressed the infertility phenotype of *wee-1.3(RNAi)* in wild type animals^31^. Five of the 44 suppressors were genes that encode subunits of the 19S RP of the 26S proteasome *(rpn-2, rpn-6.1, rpn-8, rpn-9* and *rpt-5).* The mechanism by which this suppression occurs is not currently known. Now possessing two viable 19S RP subunit mutants, we wondered whether complete genetic loss of a specific 19S RP subunit would also suppress the *wee-1.3(RNAi)* infertility. We subjected feminized *fog-2(q71), rpn-12(av93)* and *rpn-10(ana7)* mutants to either *control(RNAi)* or *wee-1.3(RNAi)* conditions. All *fog-2(q71), rpn-12(av93)* and *rpn-10(ana7)* hermaphrodites were mated to wild type males as a source of sperm during the various RNAi treatments. Mated *fog-2(q71)* females treated with *wee-1.3(RNAi)* have an average brood close to zero, while those treated with *control(RNAi)* had a large brood (Figure 4E, brood = 0.1 ± 0.05 versus brood = 96.64 ± 3.94). However, mated *rpn-12(av93)* and *rpn-10(ana7)* animals treated with *wee-1.3(RNAi)* show significant increases in the average brood size compared to *fog-2(q71)* animals (Figure 4E, brood = 11.74 ± 2.44, p value = 0.0001 and brood = 14.45±1.42, p value = 1.63e-6 respectively). To confirm that decrease in proteolytic function do not suppress WEE-1.3-depletion infertility, we also treated *control(RNAi)* or *wee-1.3(RNAi)* animals to either a control drug treatment or 100μM bortezomib. General inhibition of the proteasome with bortezomib does result in a lower brood (average brood 6.3 ± 1.483) with 47% of treated animals producing 0 brood than the control DMSO treated animals (average brood 117.208 ±2.97), however bortezomib treatment does not suppress the infertility caused by *wee-1.3(RNAi)* (Figure 4I). Combined, this data demonstrates that complete loss of RPN-12 or RPN-10 is able to suppress the *wee-1.3(RNAi)* infertility phenotype, and that this suppression is not due to the minor decrease in proteolytic activity of the proteasome within the gonads of the *rpn-12(av93)* and *rpn-10(ana7)* animals.

### Downregulation of TRA-1 partially suppresses the lack of sperm phenotype in *rpn-12(av93)* hermaphrodites

To further investigate the lack of sperm phenotype of the *rpn-12(av93)* hermaphrodites, we wanted to determine whether RPN-12 plays a previously unidentified role in the sex determination pathway. The *C. elegans* hermaphrodite germline sex determination pathway is a tightly regulated process (Figure 5A). During the L3 larval stage of hermaphrodite development, sperm production occurs via suppression of oocyte facilitating proteins (such as TRA-2 and TRA-1) by both post-transcriptional and post-translational regulators.^34–36^ Inappropriate upregulation of TRA-2 causes feminization of the hermaphrodite germ line and a complete lack of sperm.^26,36^ In *C. elegans,* the 19S RP subunit RPN-10 was shown to be involved in the sex determination pathway by specifically downregulating TRA-2 during hermaphrodite larval development and thus promoting spermatogenesis.^26^ Loss of RPN-10 increases TRA-2 expression inhibiting spermatogenesis thereby resulting in hermaphrodites with a feminized germline phenotype.^26^ Down regulation of TRA-2 or TRA-1 during larval development via RNAi in *rpn-10(lf)* mutants restores sperm production.^26^ Studies done in yeast show that RPN-12 is required for the incorporation of RPN-10 into the 19S RP of the proteasome.^20^ RNAi data in *C. elegans* show that RPN-10 and RPN-12 compensate for each other because downregulation of just one does not cause lethality, but co-depletion of both subunits causes embryonic lethality.^24^ However, our data shows that loss of RPN-12 in hermaphrodites causes a similar lack of sperm phenotype to that seen in *rpn-10* mutants indicating absence of subunit compensation during sperm production. We thus hypothesized that RPN-12 might also be playing a role in the germline sex determination pathway.

**Figure 5.**
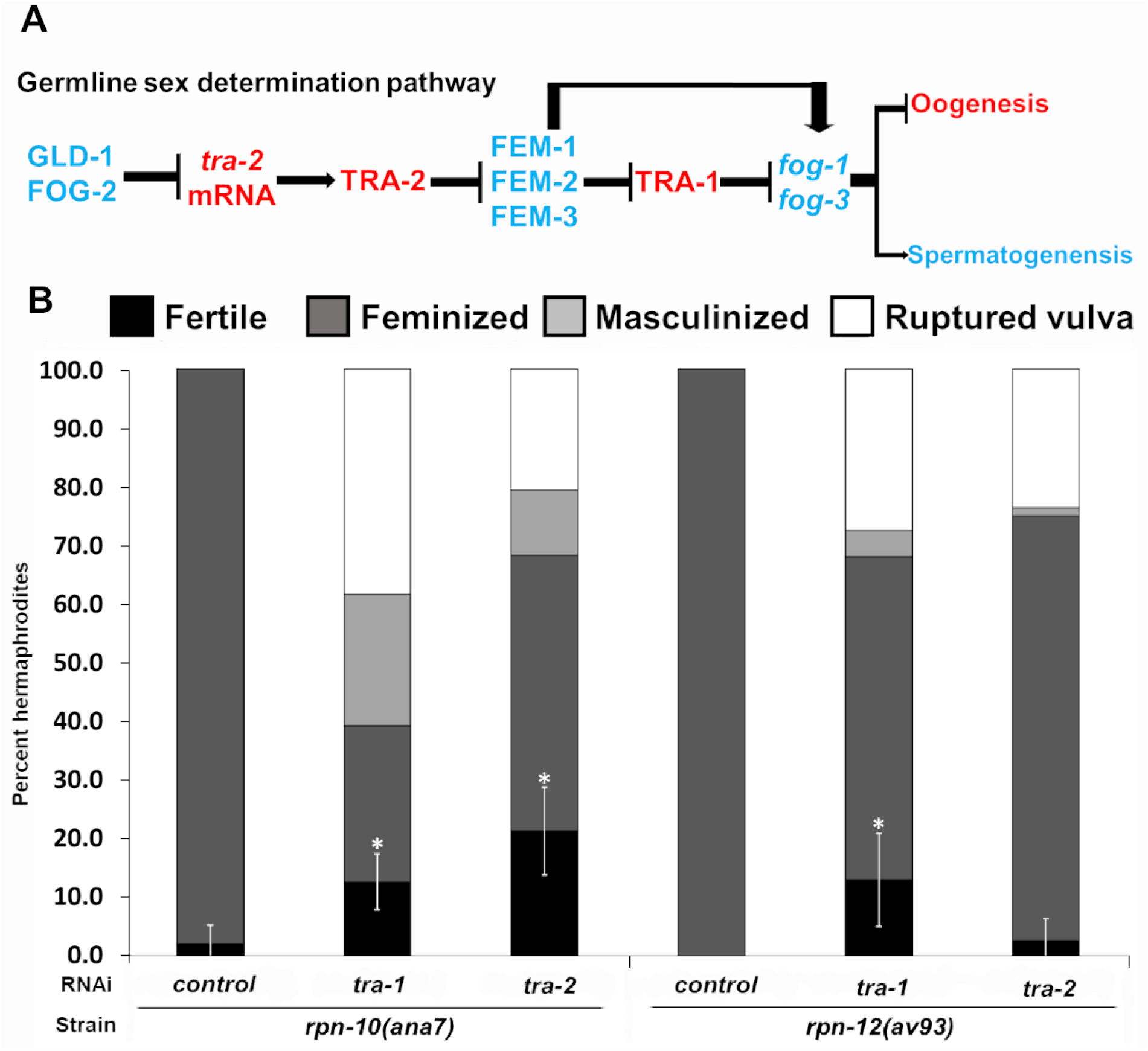
*rpn-12(av93)* hermaphrodite germline feminization phenotype can be partially suppressed by *tra-1(RNAi).* A. Schematic of the hermaphrodite germline sex determination pathway showing oogenesis facilitating proteins in red and spermatogenesis facilitating proteins in blue. Accumulation of TRA-2 and/or TRA-1 during larval development can cause germline feminization. B. Percentage of *rpn-10(ana7)* and *rpn-12(av93)* hermaphrodites exhibiting fertile (black bars), feminized (dark grey bars), masculinized (light grey bars) and ruptured through vulva (white bars) phenotypes when treated with *control(RNAi), tra-1(RNAi)* and *tra-2(RNAi).* Hermaphrodites treated from L1 stage at 25°C. All experiments were conducted at least three times and n=69-188 for each condition. * p value <0.05 determined by the Pearson’s chi-square test when compared to the mutant *control(RNAi)* data. Error bars indicate 95% confidence intervals.

To test our hypothesis, we performed *tra-1(RNAi)* or *tra-2(RNAi)* on *rpn-12(av93)* animals to see whether depletion of either TRA-1 or TRA-2 could rescue the lack of sperm production phenotype. Strong knockdown of TRA-1 and TRA-2 by RNAi during larval development can result in a masculinized germ line and some animals exhibit a ruptured vulva phenotype.^26,37–40^ Therefore, we used reduced concentrations of RNAi culture for feeding, but were still able to observe a percentage of RNAi-treated animals exhibiting masculinized germ lines and ruptured vulvas indicating our treatments are effectively downregulating TRA-1 and TRA-2 successfully. Additionally, we observed a deformed male tail phenotype upon TRA-1 and TRA-2 RNAi treatment which corresponds to another reported phenotype of depletion of TRA-1 and TRA-2, pseudomale tail morphology in XX animals (data not shown).^39,40^ We utilized our newly generated null *rpn-10(ana7)* mutant as a control. As previously reported, we saw that downregulation of TRA-1 or TRA-2 in the *rpn-10(ana7)* animals restored fertility to 12.7% or 21.3% of the treated hermaphrodites respectively when compared to the feminized *control(RNAi)* animals (Figure 5B, n=117-188, p values 0.003 and 0.000032 respectively). Depletion of TRA-1 was able to rescue the feminized phenotype of the *rpn-12(av93)* hermaphrodites (Figure 5B). We saw that upon *tra-1(RNAi),* 13% of the *rpn-12(av93)* hermaphrodites tested restored sperm production and were fertile (p value 0.001), while 55.1% still had feminized germ lines, 4.4% showed masculinization of the germ line and 27.5% were ruptured through vulva (Figure 5B, n=69). Only 2.7% of the hermaphrodites rescued sperm production (p value 0.136) in *rpn-12(av93)* animals upon *tra-2(RNAi),* while a majority (72.2%) of the animals remained feminized (Figure 5B, n=72).

Our data show that loss of RPN-12 causes minor impairment of the proteolytic activity of the proteasome in the germ line. We reason that the decreased proteolytic activity in the absence of RPN-12 during larval development causes an increase in TRA-1 levels that leads to the feminization phenotype of the *rpn-12(av93)* animals.

## Discussion

A functional proteasome is essential for the survival of all organisms. Therefore, it has been extremely challenging to elucidate any potential proteasome subunit-specific developmental functions. In this study we have characterized a viable null mutant in RPN-12, one of the 19 known subunits comprising the 19S regulatory particle of the *C. elegans* 26S proteasome. Previous studies in *C. elegans* demonstrated that knock down of RPN-12 by itself via RNAi does not cause embryonic lethality but downregulation of both RPN-12 and RPN-10 (another 19S regulatory particle subunit) together causes embryonic lethality.^24^ This implied that RPN-10 and RPN-12 are able to compensate for each other in their requirement for proper proteolytic function and viability.^24^

Our viable null single mutants for *rpn-10* and *rpn-12* confirm that these two 19S RP subunits can compensate for each other for viability, however each null mutant possesses a feminized germ line. This allows us to determine that RPN-10 and RPN-12 are both required for proper germ line development and cannot compensate for each other in this role. While both RPN-10 and RPN-12 are subunits of the 19S RP, they are radically different proteins. The RPN-10 protein has two predicted C-terminal ubiquitin-interacting motif (UIM) domains and an N-terminal VWF-A (von Willebrand factor, type A) domain.^26^ Meanwhile the only identifiable protein domain in RPN-12 is a PCI domain (Proteasome, COP9 signalosome, eukaryotic Initiation factor 3) which play roles in scaffolding protein complex subunits together and stabilizing protein-protein interactions.^28,29^ UIM domains have been demonstrated to recognize ubiquitin and be involved in targeting substrates for degradation, while VWF-A domains are commonly found in proteins involved in forming multiprotein complexes.^26,41,42^ Thus both RPN-12 and RPN-10 might be important individually for scaffolding the proteins together to form a functional 19S RP lid complex. We speculate that in the absence of both RPN-10 and RPN-12 the stability of the lid complex is compromised, thus RNAi depletion of both proteins simultaneously results in lethality.

The role of yeast Rpn10 and Rpn12 has been well characterized. When Rpn-10 is deleted in yeast, the cells are still viable and there is only a mild increase in ubiquitinated proteins observed.^43^ Biochemical and biophysical studies done in *Saccharomyces pombe* revealed that specific residues in Rpn12 are required for the proper incorporation of Rpn10, a major ubiquitin receptor, into the 26S proteasome.^20^ A temperature sensitive *Rpn12* mutant isolated in *S*. *pombe* was found to be synthetically lethal with a *Rpn10* deletion mutant.^44^ Therefore an intriguing hypothesis would be that *C. elegans* RPN-12 might also be required for the specific incorporation of RPN-10 into the proteasome. Future experiments must determine whether *C. elegans* RPN-10 and RPN-12 are capable of physically interacting.

We have demonstrated here that the *rpn-12* null mutant only exhibits minor impairment in proteolytic function in the germ line. Loss of RPN-10 in *C. elegans* and other systems does not negatively affect proteasome stability or general proteolytic activity.^26,42,45,46^ In *C. elegans* loss of RPN-10 has been shown to induce the SKN-1 mediated stress response pathway that aids in the organism’s survival via increasing the transcription of other proteasome subunit genes and genes involved in oxidative, heat stress response and autophagy.^46^ The slight decrease in the proteasome activity in the absence of RPN-12 may thus also activate the stress response pathways helping the *rpn-12(av93)* animals to survive. Further experiments need to be done to elucidate the mechanism by which *rpn-12(av93)* animals can survive in the absence of a functional RPN-12 lid subunit. Additionally, while we only determined germline proteolytic function activity, we propose that proteolysis throughout the somatic tissue must also only be slightly impaired in the absence of RPN-12 because otherwise the animals would be homozygous lethal as observed with other null mutants of proteasomal subunits.

While RPN-12 is not essential for the viability of *C. elegans,* its absence does cause defects in hermaphrodite reproduction. Our results show that RPN-12 null mutant hermaphrodites exhibit a germ line feminization phenotype and lack sperm (Figure 1B-D). Loss of RPN-12 in males however does not influence *C. elegans* male spermatogenesis or fertility (Figure 1E). This suggests that RPN-12 is specifically involved in the hermaphrodite germline sex determination pathway. During hermaphrodite larval development and germ line formation, a delicate balance between spermatogenesis promoting and oogenesis promoting factors must be maintained to ensure proper gamete production.^47^ Spermatogenesis in the hermaphrodites occurs during the late L3 to L4 stages during which oogenesis facilitating proteins, such as TRA-1 and TRA-2, are suppressed.^35,48^ TRA-2 is known to be suppressed at the transcript levels by RNA-binding proteins such as GLD-1, while TRA-1, a transcription factor, is post-translationally regulated via the CUL-2 E3 ligase and the FEM proteins that targets TRA-1 for proteasomal degradation.^49^ TRA-2 is upstream of TRA-1 and can regulate TRA-1 activity via inhibiting the FEM proteins. A previous study by Shimada *et al.* was the first to identify the proteasome subunit, RPN-10, as a potential negative regulator of TRA-2 levels during *C. elegans* spermatogenesis.^26^ In their study they concluded that RPN-10 acts as a ubiquitin receptor in the UFD pathway to select specific substrates for degradation.^26^ They also showed that downregulation of either TRA-2 or TRA-1 during larval development can partially restore spermatogenesis in *rpn-10(tm1349)* hermaphrodites.^26^ We were able to reproduce these results with a newly generated CRISPR null *rpn-10* mutant.

In contrast to the *rpn-10* mutant hermaphrodites, only larval RNAi-depletion of TRA-1 in the *rpn-12(av93)* hermaphrodites was able to significantly rescue the spermatogenesis defect. We propose that in the absence of RPN-12, TRA-1 levels will have risen thus promoting a feminized germline. Therefore, upon RNAi downregulation of TRA-1 in *rpn-12(av93)* hermaphrodites the levels of TRA-1 are reduced sufficiently to allow upregulation of spermatogenesis promoting genes at the appropriate time of germline development. *tra-2(RNAi)* is less effective in *rpn-12(av93)* mutants at rescuing the spermatogenesis defect as a high percentage of the RNAi treated *rpn-12* mutant animals still possessed feminized germ lines after RNAi treatment. Therefore, we propose that RPN-12 is working at the level of TRA-1 in the germline sex determination pathway, downstream of TRA-2. The fact that sperm production can be successfully rescued via down regulation of TRA-2 in *rpn-10* mutants and not in *rpn-12* mutants suggests different roles for those subunits in the germline sex determination pathway. These findings lead us to hypothesize that RPN-10 and RPN-12 while redundant for embryonic viability, possess distinct roles during hermaphrodite germline sex determination.

It is worthwhile to also consider the possibility that in the absence of RPN-12, TRA-2 is not suppressed appropriately and may accumulate at such high levels that we were not able to sufficiently downregulate TRA-2 via RNAi during spermatogenesis. Therefore, the germ line remains feminized in most *rpn-12(av93)* animals treated with *tra-2(RNAi).* We speculate that the reason we observe the TRA-2-depletion phenotype of a ruptured vulva in the *rpn-12* mutants is because the continual RNAi of TRA-2 from the L1 stage throughout larval development is effective to eventually reduce TRA-2 levels enough at a later development stage and affect vulval development. Finally, as the germline sex determination pathway is very complex, it is also possible that RPN-12 regulates another factor in the pathway besides TRA-1 or TRA-2. In the absence of *rpn-12* this unknown factor would exhibit misexpression and promote oogenesis. These differing potential mechanisms for RPN-12 action during germline sex determination remain to be elucidated in the future and the *rpn-12(av93)* strain therefore can be a useful tool to further genetically dissect the germline sex determination pathway.

Interesting to us, when the *rpn-12(av93)* hermaphrodites are supplied with sperm via mating with wild type males, the RPN-12 deficient oocytes can be cross-fertilized but the animals exhibit a reduced average brood compared to that of control mated animals. This suggests potential defects in oocyte quality in the *rpn-12(av93)* mutants. We demonstrate that *rpn-12(av93)* animals exhibit overall increased levels of WEE-1.3 in the oocytes, and aberrant oocyte nuclear accumulation of WEE-1.3 (Figure 4A-D). WEE-1.3 is a crucial cell cycle regulator that maintains oocyte meiotic arrest by inactivating Maturation Promoting Factor (MPF) via CDK-1 phopshorylation.^30^ RNAi depletion of WEE-1.3 starting from the L4 stage causes precocious oocyte maturation, poor quality oocytes, and infertility.^30,31^ In wild type oocytes, WEE-1.3 is localized to the cytoplasm, ER, and exhibits peak expression in the perinuclear region.^31^ We hypothesize that the aberrant WEE-1.3 expression is the reason why the *rpn-12(av93)* and *rpn-10(ana7)* animals are subfertile and results in poor quality oocytes. Future investigations will be directed towards characterizing the oocyte defects in *rpn-12(av93)* and *rpn-10(ana7)* animals that have been supplied with sperm continuously via mating.

Since loss of RPN-12 or RPN-10 does not impair proteolytic function severely in the germ line, we did not expect to see protein accumulation in the germ line such as that which we observed with overall increased nuclear WEE-1.3 expression. If the accumulation and aberrant nuclear localization of WEE-1.3 is due to compromised proteolytic activity of the proteasome and thus a decreased rate of WEE-1.3 turnover, then chemical inhibition of the proteasome with bortezomib should have also caused the same phenotype. However, we saw that treating with bortezomib only caused a cytoplasmic increase in the levels of WEE-1.3 and did not result in an increase in the nuclear accumulation of WEE-1.3 above cytoplasmic levels (Figure 4F-H). This suggests that while normal proteolytic function is required to maintain wild-type WEE-1.3 levels and protein turnover in the oocytes, RPN-12 and RPN-10 are involved in proper localization of WEE-1.3. We are actively investigating whether there is a direct interaction between WEE-1.3 and either RPN-12 or RPN-10 that might contribute to the spatial expression pattern of WEE-1.3 within the germ line. Another supporting connection between the RPN-12 and RPN-10 and the meiotic kinase WEE-1.3 is that *rpn-12(av93)* and *rpn-10(ana7)* mutants suppress the *wee-1.3(RNAi)* infertility phenotype (Figure 4E). Again, our data supports that that this is not due to a proteolytic function of RPN-12 or RPN-10 because chemical depletion of proteolytic activity via bortezomib does not suppress *wee-1.3(RNAi)* hermaphrodite infertility (Figure 4I). This supports our idea that RPN-12 may play non-canonical roles in the *C. elegans* germ line that is separate from its function as a proteasomal subunit.

In conclusion, we have demonstrated that RPN-12 is not essential for the proteolytic function of the 26S proteasome yet plays a crucial role in specific pathways that regulate reproduction, including germline sex determination. It will be interesting to test whether RPN-10 and RPN-12 interact in *C. elegans* and to determine if RPN-12 interacts with specific non-proteasomal proteins that are involved in *C. elegans* reproduction.

## Materials and methods

### Strains

All strains were maintained at 20°C.^50^ Bristol strain N2 was used as the wild type strain. Strain AG343 *rpn-12(av93)* is a complete loss of function endogenous deletion mutant.^25^ Strain JK574 *fog-2(q71)* was used as a control feminized germline strain. The germline specific indicator strain for proteasome function IT1187 *(unc-119(ed3) III; kpIs100 [pie-1p::Ub(G76V)::GFP::H2B::drp-1 3’ UTR; unc-119(+)])* was a gift from the Subramanium lab^32^. The strain IT1187 was crossed into the *rpn-12(av93)* to generate *(unc-119(ed3) III; kpls100[pie-1p::Ub(G76V)::GFP::H2B::drp-1 3’ UTR; unc-119(+)]); rpn-12(av93).*

### Strain generation

Strains in this study were generated using CRISPR/Cas9 genome editing technology following the direct delivery method developed by Paix *et al.* 2015.^51^ The Co-CRISPR method using *unc-58* or *dpy-10* was performed to screen for desired edits.^52^ Specificity of the crRNAs were determined using UCSC genome browser and http://crispr.mit.edu/. ApE plasmid editor was used for sequence analysis to select PAM sites and design primers. The edits were confirmed using PCR. At least two independent strains were generated for each edit and the resulting edited strains backcrossed at least 5 times before being utilized.

Strains WDC6 *rpn-12(ana6[gfp::rpn-12])* and WDC2 *wee-1.3(ana2[gfp::wee-1.3])* were generated by inserting Superfolder GFP sequence at the N-terminus immediately after the start ATG.^53^ Repair templates for the GFP strains were generated by PCR amplifying Superfolder GFP from pDONR221. The *gfp::rpn-12* repair template: forward primer 5’-tatcaaattaaaacattattggatttaagaaaatg<u>tccaagggagaggagctctt</u>-3’ (oAKA498) and reverse primer 5’-tcctttgcccacacagccagaagatttttatgggcagcaga<u>cttgtagagctcgtccattc</u>-3’ (oAKA499). The *gfp::wee-1.3* repair template: forward primer 5’-gcgtatgattaatttattcattttcagtgaaaatg<u>tccaagggagaggagctctt</u>-3’ (oAKA423) and reverse primer 5’-ccatttcgaatggaatccatcgaggagttaccctcggtatcatc<u>cttgtagagctcgtccattc</u>-3’ (oAKA424). Underlined region is Superfolder GFP sequence. To introduce double strand breaks near the start codon of *rpn-12* and *wee-1.3,* the crRNA34 5’-agccagaagatttttatggg-3’ *(rpn-12)* and crRNA29 5’-agccagaagatttttatggg-3’ *(wee-1.3)* (Horizon Discovery Ltd) were used with purified Cas9 (gift from Golden lab). PCR confirmation of a successful edit was performed utilizing the following primers: oAKA415 5’-accttcacagaatcgtcgag-3’ forward and oAKA506 5’-cattcaaatcgctggaggca-3’ reverse for *gfp::rpn-12* strain. oAKA67 5’-cttctctgcaccgcaaaaat-3’ forward and oAKA55 5’-gaacgagcctcttccaatga-3’ reverse for *gfp::wee-1.3* strain.

The loss of function endogenous deletion mutant strain WDC7 *[rpn-10(ana7)]* that phenocopies the classical *rpn-10(lf)* mutants was generated using two crRNAs, crRNA39 5’-acggagatttccaaccaact-3’ and crRNA40 5’-ttaggctcttcgtgtctcaa-3’ (Horizon Discovery Ltd) targeted to introduce two cuts at both ends of the gene and repaired with an ssODN 5’-ttcggaatacatgcgcaacggagatttccaaccaactcaagaggaacgagctcgtcaagccgccgccgctgct-3’ (IDT, Inc.) that creates a 987bp deletion in the coding region of *rpn-10.* PCR confirmation of a successful edit was performed utilizing the following primers: oAKA523 5’-tctcctacactgacaacacc-3’ forward and oAKA524 5’-gtggctggatggtgattcaa-3’ reverse. *gfp::wee-1.3; rpn-12(av93)* and *gfp::wee-1.3; rpn-10(ana7)* were generated by crossing *gfp::wee-1.3* to *rpn-12(av93)* and *rpn-10(ana7)* respectively.

### Fertility assays

Self-brood counts of wild type and *rpn-12(av93)* hermaphrodites were assessed by placing single L4 stage hermaphrodites on individual OP50 seeded plates, passaging parent worms to new plates every 24hrs for 72 hours and then scoring the total brood from two independent trials (n>50 in each trial). Mated brood counts were assessed by adding five wild type males to one L4 stage *rpn-12(av93)* or wild type hermaphrodite on individual plates, passaging the parent worms to new plates every 24hrs for 72hrs and then scoring the total brood from three independent trials. Male fertility was assessed by mating wild-type or *rpn-12(av93)* males to *fog-2(q71)* females by mating one age synchronized, young adult male to one young adult female on individual plates and passaging the parents to a fresh plate every 24hrs for 72hrs and scoring brood. All fertility assays were conducted at 20°C. Total brood consists of embryos and hatched larvae.

### Live imaging

10μl of anesthetic (0.1% tricane and 0.01% tetramisole in 1X M9 buffer) was added to a 3% agar pad on a slide and 10-15 live worms were transferred to the drop of anesthetic. A glass coverslip was slowly lowered to cover the samples and the coverslip edges were sealed with nail polish and allowed to dry before imaging. The images were obtained on a Nikon Ti-E-PFS inverted spinning-disk confocal microscope using a 60x 1.4NA Plan Apo Lambda objective. The microscope consists of a Yokowaga CSU-X1 spinning disk unit, a self-contained 4-line laser module (excitation at 405, 488,561, and 640nm), and an Andor iXon 897 EMCDD camera. Fluorescence intensities were quantified and image editing done using NIS-elements software.

### Whole worm DAPI staining

Whole worms were fixed with 95% ethanol on a charged microscope slide and incubated with 10mg/ml DAPI diluted 1:500 in 1X PBST. Extra dye was washed off with 1X PBST and excess buffer wicked off with a Kimwipe before covering the sample with a coverslip, drying and sealing the coverslip edges with nail polish. Samples were imaged using the microscopy conditions described previously.

### Immunofluorescence

Tube staining method was performed on dissected gonads fixed in 3% paraformaldehyde and methanol.^54^ The samples are washed using 1XPBST (0.1% tween), blocked with 30% NGS and incubated with primary antibodies at 4°C overnight. Appropriate secondary antibodies were added and incubated at room temperature for 1 – 2hrs followed by three washes with 1XPBST with DAPI included in the final wash and samples were mounted on a 3% agar pad with Vectashield mounting medium. The primary antibodies used in this study are: a mouse monoclonal proteasome 20S core particle antibody that recognizes alpha 1,2,3,4,5,6,7 subunits (1:200, Enzo Life Sciences, Cat. No. BML-PW8195) and rabbit GFP antibody (1:500, Novus Biologicals, Cat. No. NB600-308). Secondary antibodies were goat-anti-mouse Alexa Fluor 568nm and goat-anti-rabbit Alexa Fluor 488 (1:1000, Invitrogen).

### Proteasome inhibition

Bortezomib treatment was performed following the protocol published by Lehrbach et al. 2019 with minor modifications.^33^ Briefly, bortezomib (LC Laboratories, Cat. No. B-1408) was reconstituted in DMSO (Sigma, Cat. No. 2650). Bortezomib was diluted to 100μM in 1X M9 buffer and added on top of MYOB agar plates seeded with OP50 bacteria or RNAi bacteria. L4 stage larvae were added to either drug treated or DMSO treated plates and incubated at 24°C. Fertility assays or imaging was performed using treated animals as described above.

### Fluorescence intensity measurements of oocytes

All images were collected utilizing the same collection parameters. NIS-elements software was used to measure the fluorescence intensity across a −2 oocyte. A midplane Z-stack section was selected and a line drawn through the nucleus of the −2 oocyte that extended from near the distal to near the proximal oocyte plasma membrane. We ensured that sufficient cytoplasmic regions on either side of the nucleus were included in the region covered by the line. The software then calculated the intensity across the line at the various points. We manually determined the location of the perinuclear regions for each line. A line graph was then generated in Microsoft Excel to determine average intensities across the relative distances with each set of samples.

### RNA interference (RNAi) feeding

The following RNAi clones were obtained from the Open Biosystems ORF-RNAi library: *wee-1.3(Y53C12A.1)* and *smd-1(F47G4.7).* The *tra-2(C15F1.3)* RNAi clone was obtained from the Ahringer RNAi library and *tra-1(Y47D3A.6)* RNAi clone from the Vidal ORFeome RNAi library.^55^ The *smd-1(RNAi)* was used as the control RNAi due to it having no reported phenotype yet being able to trigger an RNAi response. RNAi depletion of WEE-1.3 was performed on *rpn-12(av93)* and *fog-2(q71)* hermaphrodites at 20°C. Sperm were introduced by mating both strains to wild type males. On Day 1, 10-20 L4 larvae hermaphrodites of *rpn-12(av93)* or *fog-2(q71)* along with approximately 30 wild type males were placed on RNAi plates [MYOB containing 1mM IPTG (GoldBio, Cat. No. I2481C) and seeded with the appropriate dsRNA expressing HT115 cells]. On Day 2, the hermaphrodites were singled out to individual plates with 10 fresh wild type males each. On Day 3, the parent worms were removed and embryos were allowed to develop for 24hrs before the brood was scored on Day 4. For the TRA suppression experiments: *smd-1, tra-1* and *tra-2* RNAi feeding was performed on *rpn-12(av93)* and *rpn-10(ana7)* at 25°C starting from the L1 stage. L1 larvae were obtained by sodium hypochlorite treatment of gravid animals to age synchronize the worms. Upon reaching the L4 stage the hermaphrodites were transferred to new RNAi plates. After 24hrs the worms were imaged to score the number of hermaphrodites that had embryos in their uteri (fertile), had no embryos in the uteri (feminized), contained only sperm and no oocytes (masculinized), or were dead on the plate with intestines protruding out (ruptured vulva). Chi-square test was performed to determine the statistical significance of percent hermaphrodites with various phenotypes between treatment groups.

## Acknowledgements

We thank the Duttaroy and Robinson Labs for sharing equipment and reagents; Dr. Andy Golden, Dr. Katherine McJunkin and Dr. Kuppuswamy Subramaniam for kindly providing the pDONR221, *tra-1(RNAi)* clone, and IT1187 strain respectively; and members of the Baltimore Worm Club for helpful discussions. We thank the following people for comments on the article: Andy Golden and members of the Allen lab. This research was funded by a Department of Defense grant to AKA (#W911NF1810465). LMF was supported both by the DoD grant and the Howard University Graduate School Just-Julian Fellowship. The spinning disk confocal microscope was acquired through a Department of Defense HBCU/MI Equipment/Instrumentation Grant (#64684-RT-REP) to AKA. Some strains were provided by the CGC, which is funded by NIH Office of Research Infrastructure Programs (PO OD10440). Author contributions: LMF designed the study, conducted the experiments, interpreted data, and wrote the manuscript. JE conducted experiments and interpreted data. AKA designed the study, interpreted data, and wrote the manuscript.

## Notes

### Competing Interest Statement

The authors have declared no competing interest.

